# Polysomes and mRNA control the biophysical properties of the eukaryotic cytoplasm

**DOI:** 10.1101/2024.11.14.623620

**Authors:** Vamshidhar R. Gade, Stephanie Heinrich, Matteo Paloni, Pablo A. Gómez-García, Ajla Dzanko, Alexandra Oswald, Désirée Marchand, Sarah Khawaja, Alessandro Barducci, Karsten Weis

## Abstract

The organization and biophysical properties of the cytoplasm influence all cellular reactions, including molecular interactions and the mobility of biomolecules. It has become clear that the cytoplasm does not behave like a simple fluid but instead is a densely crowded and highly organized environment. Yet, the detailed properties of the cytoplasm, the molecular mechanisms that control them and how they influence the biochemistry of cells remain poorly understood. Here, we investigate the diffusive properties of the cytoplasm *in silico* and *in vivo,* employing mRNPs (<u>m</u>essenger <u>r</u>ibo<u>n</u>ucleo<u>p</u>rotein) and GEM (<u>g</u>enetically <u>e</u>ncoded <u>m</u>ultimeric) particles as rheological probes in proliferating cells. We demonstrate that cytoplasmic diffusivity increases upon polysome disassembly due to translation inhibition or upon a reduction in mRNA levels. Reducing ribosome concentration by up to 20-25% without a change in polysome levels has no effect *in vivo*. In addition, we show that upon polysome disassembly, mRNA condensation into P-bodies does not affect cytosolic diffusion in budding yeast. Altogether, our results show that mRNAs and their organization into polysomes control the biophysical properties of the eukaryotic cytoplasm.

**Highlights:**

- Polysomes control the biophysical properties of cytoplasm.
- mRNP and GEM mobility is enhanced upon translation inhibition that leads to polysome disassembly
- Perturbation of mRNA levels leads to an increase in cytosolic diffusion.

## Introduction

The eukaryotic cytoplasm is extremely dynamic and undergoes continuous reorganization. Different macromolecules (RNA, proteins) and membrane-bound or membrane-less organelles are densely packed in a confined space, crowding the intracellular environment. Crowding is further influenced by the concentration of ions and osmolytes that control water fluxes and cell size^1,2,3^. In the past decades, it has become evident that the densely packed cytoplasm is not a uniform solvent like water, but rather a heterogeneous viscous fluid^4,5^ in which molecules cannot move freely and instead undergo anomalous diffusion. Using various techniques^6–9^, the biophysical nature of the cytoplasm has been described as either porous, viscoelastic, glassy or heterogenous, yet the detailed properties of the cytoplasm are not well understood. Furthermore, it is poorly characterized how the biochemistry of cells is affected and regulated by this confined and complex environment to ensure cellular homeostasis and survival.

The crowded nature of the cytoplasm can hinder the motion of biomolecules, but it can also enhance the interactions of macromolecules due to depletion attraction forces^4,10^. In addition, macromolecular crowding affects biochemical processes such as protein folding, aggregation, and transport in the cytoplasm^11,12^. The impact of crowding significantly depends on the relative size and concentration of the crowder and the affected biomolecules. For instance, the diffusion of small macromolecules might not be influenced by changes in the concentration of crowders that are significantly larger in size^4,5^.

Since one strong effect of macromolecular crowding is manifested in the intracellular diffusion rate of biomolecules, passive rheology approaches have been successfully used to analyze the biophysical properties (diffusion, motility and viscosity) of the cytoplasm and nucleus of cells^13,14^. For example, single-particle tracking of endogenous molecules such as transcription factors has demonstrated their dynamics and exploratory nature in the nucleus^15,16^. Similarly, tracking of single mRNPs has provided information on their spatiotemporal distribution and the kinetics of movement in different compartments of the cell^17,18–20^. In addition, several exogenous rheological probes have been developed to investigate the diffusive properties of the cell via single particle tracking and high acquisition rate microscopy^13,14^. Intriguingly, these passive rheology approaches have revealed that cells not only control physiological processes such as their gene expression patterns but also regulate the biophysical properties of the cytoplasm following exposure to stress conditions^21–23^, during different cell cycle stages^24–26^ or in response to changes in cell size^27^. For instance, we and others have shown that in bacterial, budding and fission yeast cells the intracellular diffusion of macromolecules is drastically altered upon starvation and entry into dormancy^21–23,28,29^. Using the aforementioned passive rheology approaches, it was shown that the cytoplasm transitions from a liquid to a solid-like state by modulating cell volume, crowding and intracellular pH^21–23^. Similarly, in mammalian cells, intracellular diffusion significantly increases upon cell cycle transitions^24^ or entry into senescence^27^.

Mechanistically, changes in diffusion rates can be controlled by tuning the concentration of macromolecules in the cytoplasm, e.g. by adjusting the cell volume^21^. Large membrane-enclosed organelles can also affect the intracellular diffusion of biomolecules, as can mitochondria and their dynamic remodelling. At the mesoscale level (20-200 nm) it was proposed that ribosomes (∼30 nm), which occupy around 20% of cellular volume, act as major crowders in the cytoplasm^7^. This was demonstrated by inhibition of TOR1 with rapamycin or deletion of *SFP1* to reduce ribosome biogenesis and ribosome number^7^. However, inhibition of TOR1 and deletion of *SFP1* has many consequences on cellular physiology and, among other effects, results in a significant decrease in translation levels and the number of polysomes in the cytoplasm^30,31^. Since cytoplasmic ribosomes do not predominantly exist as independent entities but are mostly actively engaged in translation, this raises a fundamental question: does the translational status of a cell determine the macromolecular crowding in the cytoplasm? Since the translation status in cells is controlled by both ribosomal amounts and mRNA levels, it also remains uninvestigated whether mRNA levels, size distribution or subcellular localization control the biophysical properties of the cytoplasm.

Here, we use messenger ribonucleoprotein particles (mRNPs) and two sizes of genetically encoded multimeric particles (GEMs)^7^ as nano-rheological probes to investigate intracellular crowding and diffusion. We demonstrate that disassembling polysomes results in a significant increase in intracellular diffusion in proliferating yeast and mammalian cells. Depleting mRNA levels in the cytoplasm by perturbing nuclear mRNA export also leads to an increase in intracellular diffusion. In silico approaches confirm that the higher-order organization of ribosomes into polysomes critically affects intracellular diffusion rates. Altogether we propose that translational status and mRNA levels organize the eukaryotic cytoplasm and control its biophysical properties.

## Results

### Translational inhibition that results in polysome disassembly enhances mRNP mobility in the yeast cytoplasm

To determine how translation impacts the biophysical properties of the cytoplasm, we examined the mobility of endogenous mRNPs in different translation states by tracking single mRNAs in yeast. Since it is unclear whether the number of RNA-binding proteins (RBPs) scales stoichiometrically with the length of the mRNA^32–34^, we first asked how mRNA length affects the mobility of mRNPs in the subcellular environment. To address this question, we tagged three different endogenous mRNAs (*GFA1, TRA1* and *MDN1*) in a length range from 2.1 kb (*GFA1)* to 14.7 kb (*MDN1*, the longest mRNA in yeast) with a bacteriophage stem-loop system (PP7) at the 3’ end for visualization^35^ (Figure 1A). We imaged these three mRNAs of different lengths using a previously reported custom-built fast-acquisition microscope to image single mRNAs at ∼16ms/frame^29^ and performed a mean-squared displacement (MSD) analysis. Surprisingly, only subtle differences in the mobility of the three mRNAs could be observed in the MSD curves (Figure 1B). We also determined the diffusion coefficient and found no significant differences in the apparent diffusion coefficients (D_ensemble_) of the three mRNAs, indicating that an increase in mRNA length does not significantly slow down intracellular mRNA diffusion. Our MSD analyses suggested that individual mRNAs can undergo different motion types and display differences in the anomalous exponent α, indicating a sub-diffusive movement. We therefore asked if mRNAs of different lengths display different motion types or show differences in their confinement in the cytoplasm. Since the distribution of the diffusion coefficient of the individual trajectories is high (Figure S1A), we grouped all trajectories into two motion types, which we termed ‘confined’ with α < 0.8, and ‘non-confined’ with α > 0.8. As shown in Figure 1D, no significant difference in the motion type of the three mRNAs could be observed. In conclusion, our results suggest that the mobility of the mRNAs in the subcellular environment is similar, despite having >5-fold difference in length.

**Figure 1:**
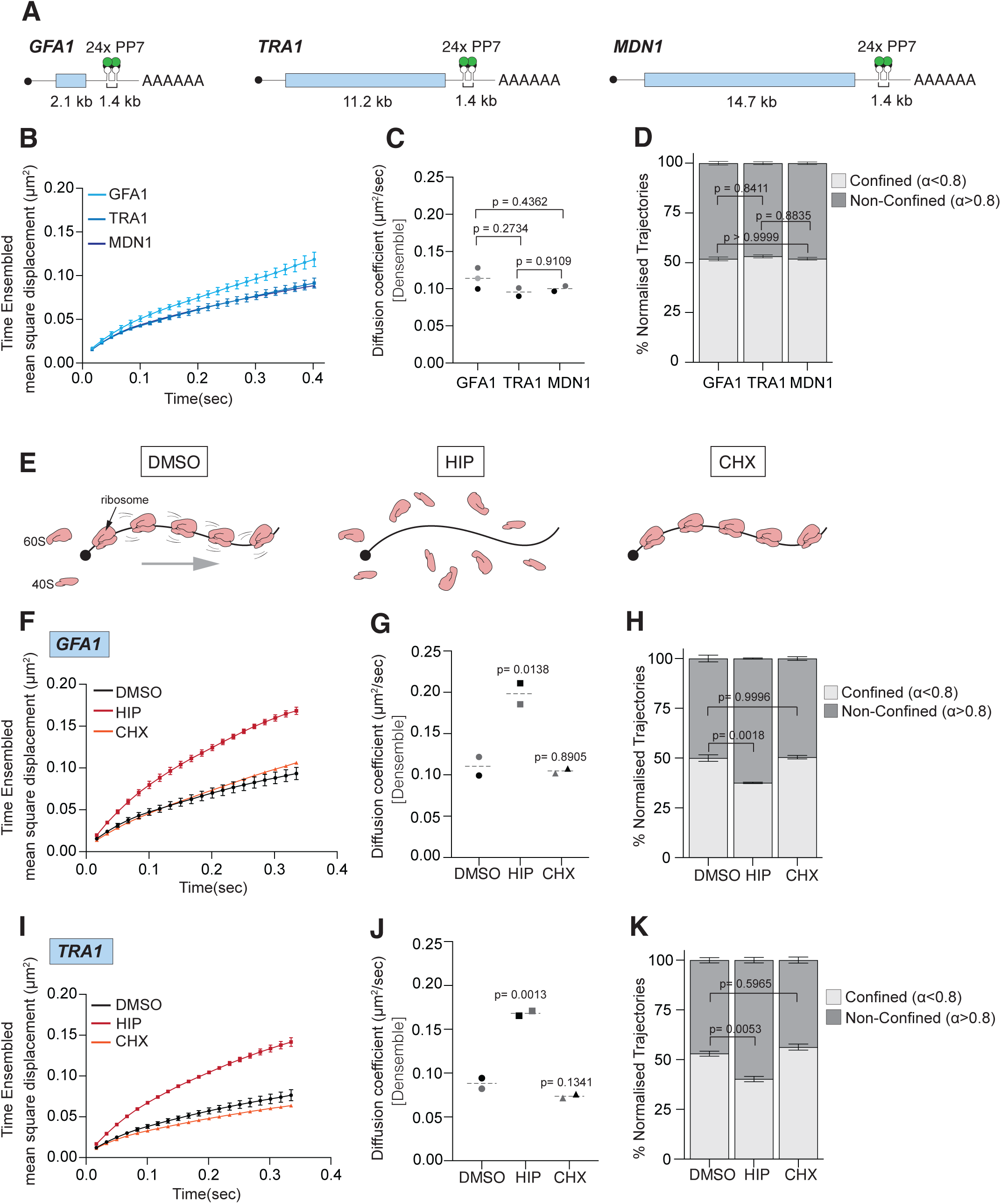
Inhibition of translation that leads to polysome disassembly increases mRNP dynamics within the cytoplasm. A) Schematic of PP7 tagging of *GFA1, TRA1* and *MDN1* mRNAs. Twenty-four PP7 stem-loops are integrated at the 3’ end after the STOP codon. GFP-tagged PP7 coat protein (PP7CP-GFP) binds to PP7 stem loops with high affinity allowing single-RNA visualization. B) Time ensembled averaged mean square displacement of three mRNAs (*GFA1, TRA1* and *MDN1*). Shown are means±SEM of two (*TRA1, MDN1*) or three (*GFA1*) biological replicates performed on different days. C) Median diffusion coefficient of three mRNAs (*GFA1, TRA1* and *MDN1*) from two (*TRA1, MDN1*) or three (*GFA1*) biological replicates performed on different days (median±SEM). Statistical comparison was performed with ordinary one-way ANOVA using Tukey’s multiple comparison test. Non-significant p value >0.05. Number of tracks (n) for GFA1 (n = 18165), TRA1 (n = 3968), MDN1 (n = 44945). D) Confined and non-confined trajectories of two (*TRA1, MDN1*) or three (*GFA1*) biological replicates performed on different days (median±SEM). Statistical comparison was performed with two-way ANOVA with Tukey’s multiple comparison test. Non-significant p value >0.05. E) Schematic illustration of translation inhibition with hippuristanol (HIP) and cycloheximide (CHX) for polysome disassembly and stabilization. F&I) Time ensembled averaged mean square displacement of *GFA1* and *TRA1* mRNA upon translation inhibition with HIP and CHX from two biological replicates performed on different days. Shown are means±SEM. G&J) Median diffusion coefficient of GFA1 and TRA1 mRNA upon translation inhibition with HIP and CHX from two biological replicates performed on different days. Statistical comparison was performed with ordinary one-way ANOVA using Dunnett’s multiple comparison test. Non-significant p value >0.05, significant p value <0.05. H&K) Polysome disassembly decreases confined trajectories of *GFA1* and *TRA1* mRNA upon HIP treatment. Statistical comparison was performed with ordinary one-way ANOVA with Dunnett’s multiple comparison test. Non-significant p value >0.05.

As most cytoplasmic mRNAs are actively engaged in translation and are bound by multiple ribosomes^36^, we next investigated whether perturbing translation affects mRNP mobility. To test this, we focused on *GFA1* and *TRA1* mRNAs where the difference in length is around 5-fold. We inhibited translation either with (a) the eIF4A inhibitor hippuristanol^37,38–40^ (HIP), which blocks initiation and results in ribosome run-off and polysome collapse by relieving the mRNAs of ribosomes, or (b) the translation elongation inhibitor cycloheximide (CHX), where ribosomes are blocked in the elongation cycle and remain stuck on the mRNA (Figure 1E). Upon HIP treatment, the mobility of both *GFA1* and *TRA1* mRNA was enhanced (Figure 1F&1I) and the diffusion coefficient (D_ensemble_) significantly increased (Figure 1G&J) (Figure S1A&C). Furthermore, a significant reduction in the percentage of the confined trajectories (Figure 1H&K) and an increase in the confinement radius (Figure S1B&D) upon HIP treatment was detected. This was in contrast to cycloheximide treatment, where no change in the diffusion could be observed. Taken together, our results suggest that the diffusion of endogenous mRNA molecules appears to be independent of their length but is significantly dependent on their association with ribosomes and their organization into polysomes.

### Polysomes control mesoscale biophysical properties of the cytoplasm in budding yeast

It was previously reported that ribosomes are the major crowder in the cytoplasm at the mesoscale level (20-200 nm)^7,41^. Because the majority of ribosomes in the cytoplasm are organized into polysomes by translating mRNAs^36^ and given the observed effects of polysomes on mRNP diffusion, we hypothesized that polysomes might be the effective crowder in the cytoplasm of proliferating cells. To test this hypothesis, we expressed 40 nm GEM particles in yeast and performed live cell imaging using highly inclined and laminated optical sheet (HILO) microscopy. This enabled us to image these particles at 100 frames/sec. We collected thousands of individual tracks (∼13,000) to extract the diffusion coefficient of 40 nm GEMs. Consistent with a previous report in *S. pombe*^8^, we observed significant heterogeneity in the diffusivity of the particles (Figure S2A), reflecting the non-uniform nature of the cytosolic environment. After an acute treatment of ∼30 min with HIP (10 μM) to induce polysome disassembly, the mobility of 40 nm GEMs increased, as shown by the MSD curves (Figure 2A). In contrast, treating cells with CHX resulted in no change in the mobility of 40 nm GEMs (Figure 2A). Similarly, the median diffusion coefficient (D_ensemble_) was significantly higher upon HIP treatment (Figure 2B) but was not affected upon treatment with CHX (Figure 2B).

**Figure 2:**
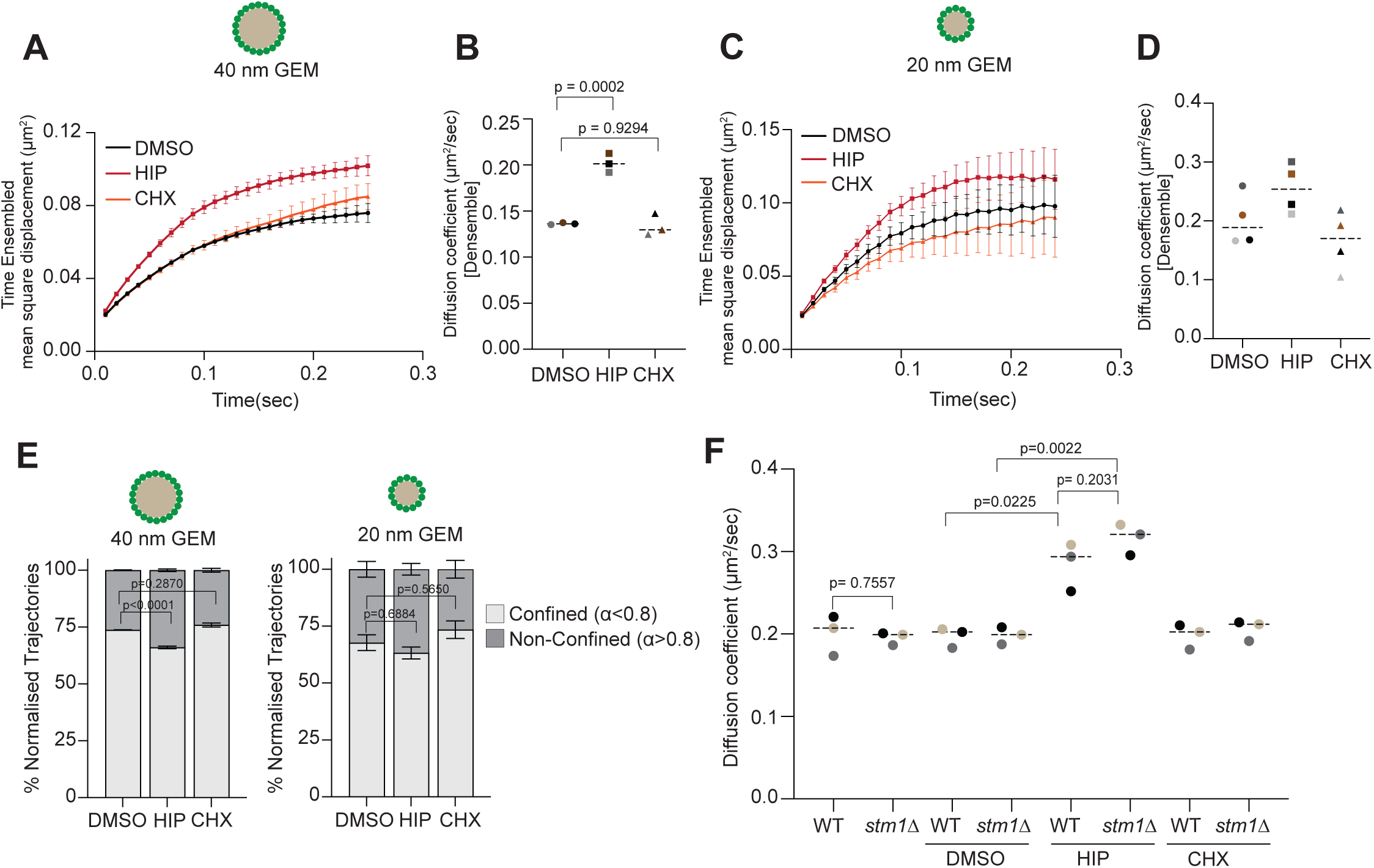
Polysome disassembly enhances cytosolic diffusion in budding yeast. A) Time ensembled averaged mean square displacement of 40 nm GEMs treated with HIP and CHX. Shown are means±SEM of three biological replicates performed on different days. B) Median diffusion coefficient of 40 nm GEMs upon treatment with HIP and CHX from three biological replicates performed on different days (median±SEM). Statistical comparison was performed using Welch’s t-test. Non-significant p value >0.05, significant p value <0.05. C) Time ensembled averaged mean square displacement of 20 nm GEMs treated with HIP and CHX. Shown are means±SEM of four biological replicates performed on different days. D) Median diffusion coefficient of 20 nm GEMs upon treatment with HIP and CHX from four biological replicates performed on different days (median±SEM). E) Polysome disassembly decreases confined trajectories of 40 and 20 nm GEMs upon HIP treatment. Confined and non-confined trajectories shown here are data from three and four biological replicates respectively and error bars are means±SEM. Statistical comparison was performed with two-way ANOVA with Šidák’s multiple comparison test. Non-significant p value >0.05, significant p value <0.05. F) Median diffusion coefficient of 40 nm GEMs upon treatment with HIP and CHX from three biological replicates performed on different days (median±SEM) in wildtype and s*tm1*Δ. Statistical comparison was performed using Welch’s t-test. Non-significant p value >0.05, significant p value <0.05.

We next asked whether changes in the state of polysomes impact the confinement of particles in the cytoplasm and result in a change in their motion. We estimated the anomalous exponent, α, from the MSD curves and like in our mRNP analysis, classified our trajectories into two groups: confined (α < 0.8) and non-confined (α > 0.8). Based on this classification, about 70% (73.69% ± 0.23%) of the generated trajectories displayed a ‘confined motion’ in the yeast cytoplasm. Upon polysome disassembly with HIP, we noticed a reduction (66.02% ± 0.98%) in the percentage of trajectories that fall into this group, in contrast to cycloheximide treatment where no difference was observed (Figure 2E). From the confined trajectories, we further derived the distance explored by the particles given by the confinement radius. Upon polysome disassembly with HIP, the confinement radius increased in comparison to CHX treatment and untreated controls (Figure S2B). These findings show that 40 nm GEMs become less confined upon polysome disassembly.

The effect of crowding critically depends on the relationship between the size of the crowder and the affected macromolecule^4^. Since polysomes are very large complexes (estimated size range 220-250 nm) we tested whether the presence of polysomes also affects the mobility of smaller particles like 20 nm GEMs. To address this, we treated budding yeast cells expressing 20 nm GEMs with HIP and CHX (Figure 2C-E). Consistent with our experiments with 40 nm GEMs, we observed an increase in particle mobility and a concomitant increase in the median diffusion coefficient (D_ensemble_) upon HIP-induced polysome disassembly (Figure 2C&D). HIP treatment also led to a subtle decrease in the percentage of confined trajectories (Figure 2E) and an increase in the confinement radius of 20 nm GEMs (Figure S2C&D), although the magnitude of the effect was less substantial in comparison to 40 nm GEMs. Upon cycloheximide treatment, there was a marginal increase in the percentage of confined trajectories (Figure 2E) and a decrease in confinement radius (Figure S2D), Altogether, our results show that polysomes act as a crowder in the cytoplasm at the mesoscale level.

It was previously concluded that reducing ribosome number by inhibiting ribosome biogenesis affects cellular diffusion^7^. However, as discussed above, perturbation of ribosome biogenesis also impacts the polysome levels^30^ and translation rate^31,42^ of cells. Since our results demonstrate that polysomes affect diffusion, we thus sought to lower the number of ribosomes without affecting global cellular translation. For this, we targeted the ribosome preservation factor Stm1^43,44^. Using cryo-electron microscopy, it was recently reported that *STM1* deletion leads to a ∼20-25 % reduction in ribosome levels without significantly affecting polysomes and the translational capacity of cells^45,46^. Therefore, we examined whether the mobility of 40 nm GEMs changes upon *STM1* deletion Intriguingly, deletion of *STM1* has no observable impact on the median diffusion coefficient (D_ensemble_) of 40 nm GEMS compared to wild-type cells (Figure 2F). This suggests that a ∼20-25% reduction of ribosome levels does not affect cytosolic diffusion. When we disassembled polysomes in *stm1*Δ cells with HIP, we still observed an increase in diffusion. Interestingly, in the presence of HIP, the diffusion increases in *stm1*Δ cells slightly exceeded that of wild-type cells, but not significantly so (Figure 2F). These results support the notion that translating polysomes and not ribosome levels (up to 20-25%) alone control cytosolic diffusion.

### Brownian dynamics simulations predict that polysome organization modulates biophysical properties of the cytoplasm

To better understand how polysomes affect diffusion in the cytoplasm and to examine the size scale across which polysomes can act as effective crowders, we performed Brownian dynamics simulations using a simple model of polysome organization. In our simulations, we modelled ribosomes, either organized in bead-spring polymers that represent polysomes or as free particles to mimic polysome disassembly, using estimates of mRNA length distribution and average polysome size in yeast as a guide^36,47,48^. We then probed the diffusion of different-sized particles representing a range of cytoplasmic biomolecules (Figure 3A&B), considering excluded volume interactions (Figure S3B). Taking as a reference the size of an 80S ribosome (σ=1), we estimated the diffusion coefficient of particles in a range of sizes between 0.01 times (σ=0.01) and 4 times (σ=4) that of the ribosome (Figure 3C). We observed that for particles larger than half the size of the ribosome, the change in diffusion increases with the size of the probe particle upon polysome disassembly whereas for particles smaller than half the size of the ribosome the diffusion coefficient was not affected in the absence of polysomes. Correlating this with the size of our rheological probes, the effect of polysome dissolution is more pronounced for 40 nm GEMs (1.3-1.6 times the size of ribosomes) than for 20 nm GEMs (0.6-0.8 times the size of ribosomes), in agreement with our diffusion measurements in yeast. Thus, despite the simplicity of our simulations, which do not consider the complexity of factors that could affect the diffusion of biomolecules in cells, our simulations recapitulate the diffusive behavior of GEMs, and our results indicate that polysome disassembly enhances the mobility of biomolecules mostly in the mesoscale range (20-200 nm). Furthermore, this agrees with our experimental evidence that mRNPs in a size range of 50-150 nm^49–51^ also display an increase in diffusion upon polysome disassembly.

**Figure 3:**
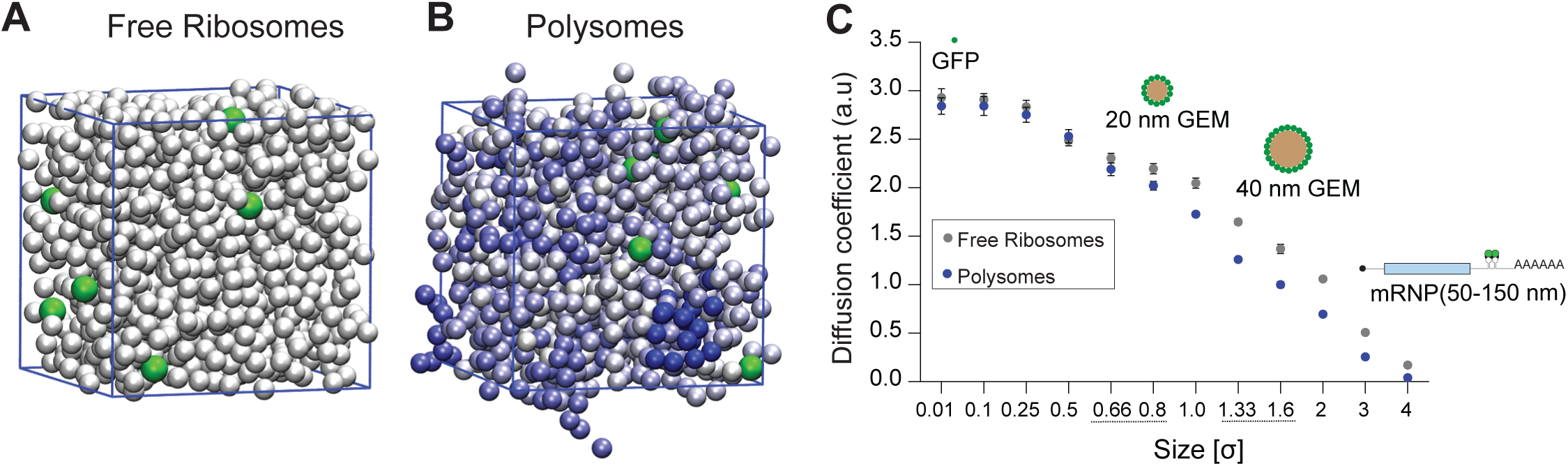
Simulations predict polysome organization controls mesoscale biophysical properties of the cytoplasm. A, B) Representative snapshots from Brownian dynamics simulations from a system mimicking polysome dissolution, where only unbound ribosomes (white) and probe particles (green) are present (A), and a system with formed polysomes (tones of blue indicate polysome length) and probe particles (green) (B). (C) Diffusion coefficients of probe particles as a function of the size of the probes in systems mimicking dissolved (green points) and formed polysomes (blue points). Sizes of the probes are relative to the size of the ribosome, set equal to 1. Values of diffusion coefficients are average values over 10 independent simulations, error bars indicate SEM.

We also used our simple model to test the effect of *STM1* deletion on the diffusion of probes of sizes in the range of 0.6-1.6 times the size of ribosomes. In our simulations, *STM1* deletion is mimicked by lowering the concentration of ribosomes in the system by 25%. Unlike in cells, this induced an increase in diffusion, however for particles in the size of 40 nm GEMs, the change is smaller than the increase due to polysome dissolution (Figure S2E). For smaller particles, similar in size to 20 nm GEMs, the effect is reduced, due to the smaller impact of the polysome state on the diffusion of the probe particles. The larger effect of ribosome concentration observed in our simulations compared to in vivo experiments is likely due to a limitation of our model in capturing the general properties of the system upon *STM1* deletion. Notably, since RNA molecules are not explicitly included in our model, a decrease of the ribosome concentration in our model corresponds to an equal decrease of the density of the system, while *in vivo* the concentration of RNA molecules and the density of the cytoplasm upon mutation are likely to remain constant, resulting in a smaller effect of the mutation.

### Reduction in cytoplasmic mRNA levels leads to increased cytosolic diffusivity

Our *in vivo* experiments and molecular simulations revealed that polysomes are significant crowders that affect macromolecular diffusion at the mesoscale above 20 nm. Since polysomes are organized by mRNAs in the cytoplasm, changes in mRNA levels are therefore expected to affect the rheological properties of the cell. To test this hypothesis, we lowered the mRNA concentration in the cytoplasm by depleting Dbp5, an essential DEAD-box helicase that mediates mRNA export ^52–54^. Dbp5 interacts with mRNPs at the cytoplasmic face of the NPC and remodels them to ensure directionality as they enter the cytoplasm^52,55^. Using FISH, we recently showed that acute depletion of Dbp5 for 2 hours using an auxin-inducible degron (AID) system leads to nuclear accumulation of bulk mRNA and a subsequent reduction in cytoplasmic mRNA levels^29^. In agreement with our hypothesis, a 2-hour depletion of Dbp5 with this AID system (Figure 4A) resulted in a significant increase in the mobility of 40 nm GEMs (Figure 4B) resulting in a pronounced increase in the median diffusion coefficient of 40 nm GEMs at both track-wise (Figure S4A) and average fits (D_ensemble_) (Figure 4C). Similar to the treatment with HIP (Figure 2E), we observed a decrease in the number of trajectories that we classified as confined (α < 0.8) upon depletion of Dbp5 (Figure 4D). Mutations in *dbp5* were previously also associated with defects in the nuclear export of pre-ribosomal subunits^56^. To ensure that the increase in the diffusion of 40 nm GEMs was not a consequence of a change in the ribosomal amounts, we investigated whether ribosomal protein levels change upon acute depletion of Dbp5. As shown in Figure S4B, no differences in the protein levels of either large (Rpl35A) or small (Rpl5A) ribosomal subunits were observed after Dbp5 depletion for 2h. Together, this corroborates our results that mRNAs via their organization into polysomes affect the rheology of the cytoplasm.

**Figure 4:**
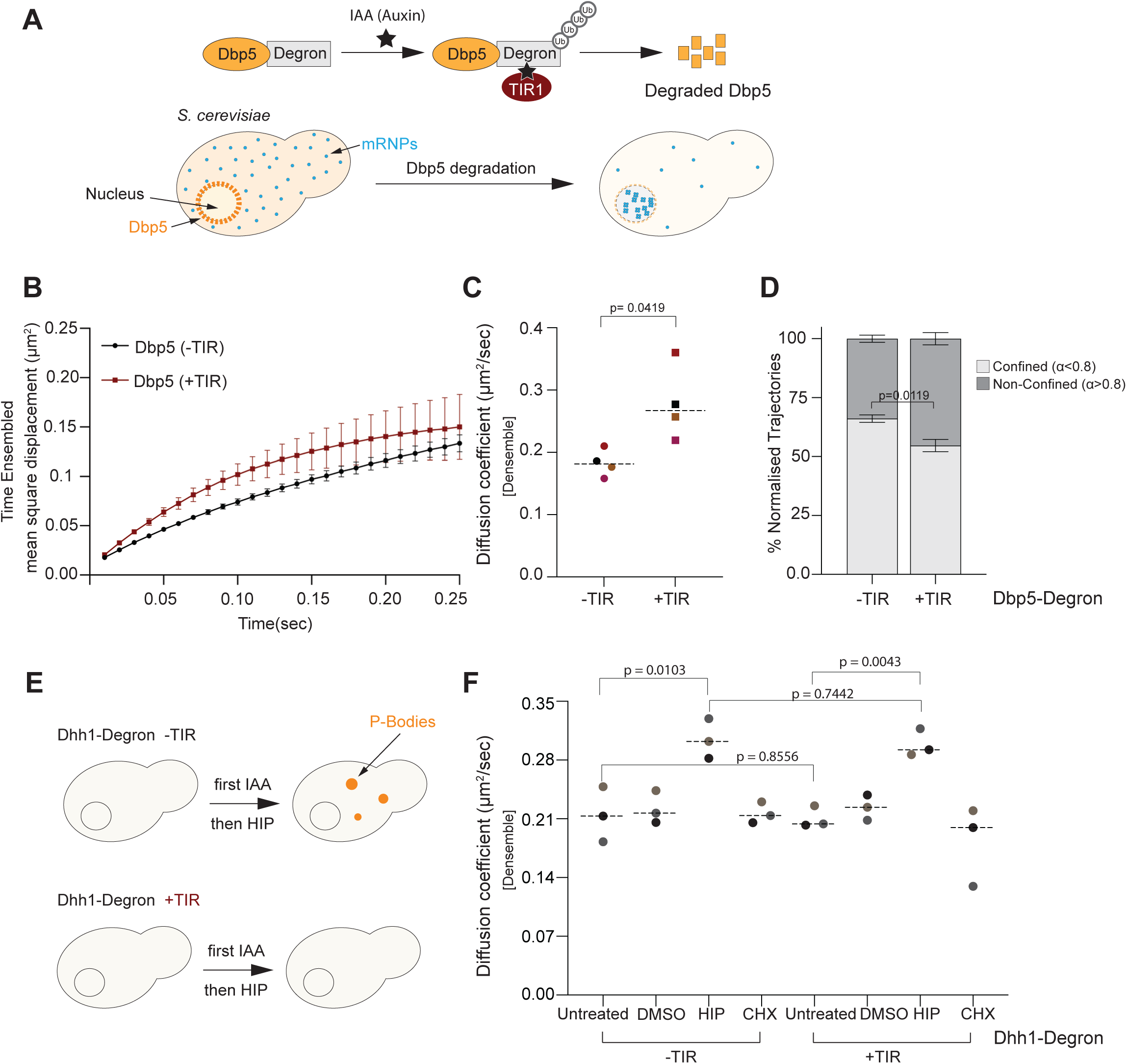
Perturbation in mRNA level enhances diffusion of 40 nm GEMs in budding yeast. A) Schematic illustration of Dbp5 depletion, which leads to mRNA sequestration in the nucleus. B) Time ensembled averaged mean square displacement of 40 nm GEMs upon acute depletion of Dbp5. Shown are means±SEM of four biological replicates performed on different days. C) Median diffusion coefficient of 40 nm GEMs of four biological replicates performed on different days. Error bars shown here are median±SEM. Statistical comparison was performed using Welch’s t-test, non-significant p value >0.05, significant p value <0.05. D) Dbp5 depletion significantly reduces confined trajectories generated by 40 nm GEMs. Data shown here is from four biological replicates performed on four different days. Error bars shown here are means±SEM. Statistical comparison was performed with 2-way ANOVA with Tukey’s multiple comparison test. Non-significant p value >0.05, significant p value <0.05. E) Schematic illustration of Dhh1 depletion and P-body formation upon HIP treatment. F) Dhh1 acute depletion (60 min) does not change the diffusion coefficient of 40 nm GEMs upon polysome disassembly. Median diffusion coefficient of 40 nm GEMs upon depletion of Dhh1 and polysome disassembly with HIP and CHX from three biological replicates performed on different days (median±SEM). Statistical comparison was performed using Welch’s t-test. Non-significant p value >0.05, significant p value <0.05.

We previously showed that HIP treatment induces the formation of P-bodies in a concentration-dependent manner and that mRNAs localize into these P-bodies (Figure 4E)^57^. This led us to test if mRNA condensation or sequestration into P-bodies contributes to the enhanced cytosolic diffusion upon HIP treatment. To address this, we first depleted the DEAD-box ATPase Dhh1 (Figure S4C), which is essential for P-body formation in budding yeast^58,59^, and then induced polysome disassembly with HIP (30 min)^58^ (Figure S4C). No significant difference in the median diffusion coefficient (D_ensemble_) of 40 nm GEMs could be observed upon Dhh1 depletion (Figure 4F) suggesting that mRNA condensation or P-body formation upon polysome disassembly is not essential for controlling intracellular diffusion in budding yeast. Therefore, we can conclude that changes in mRNA levels impact the rheological properties of the cytoplasm.

### Mesoscale cytosolic diffusion in mammalian cells is augmented by translation inhibition-induced polysome disassembly

Our results demonstrate that polysomes determine the biophysical properties of the cytoplasm in yeast at the mesoscale level. We wanted to test whether such control of intracellular diffusion is conserved and can also be seen in mammalian cells. To ensure a high acquisition rate and low phototoxicity we imaged HeLa cells expressing 40 nm GEMs using lattice light sheet microscopy. This orthogonal illumination approach allowed us to acquire images at a high frame rate (∼119 frames/sec). Consistent with our findings in yeast, a 30-minute HIP treatment led to an increase in the median diffusion coefficient (D_ensemble cell_) of 40 nm GEMs (Figure S5A&S5B). Similarly, acute treatment with puromycin for 30 min, which causes premature translation termination and polysome disassembly (Figure 5A), enhanced the mobility of 40 nm GEMs (Figure 5B) This was in contrast to CHX treatment that did not affect the MSD. The median diffusion coefficient (D_ensemble cell_) of GEMs was also significantly higher upon puromycin treatment in comparison to CHX (Figure 5C). Like in yeast, we observed significant intercellular heterogeneity in the diffusion coefficients generated by GEMs (Figure 5C&S5C). We observed that around 50% of the trajectories are confined in mammalian cells (α < 0.8; Figure 5D). In contrast to yeast, polysome disassembly did not result in any significant change (p = 0.3640) in the percentage of confined trajectories in mammalian cells (Figure 5D). However, we noticed that the median confinement radius derived from the confined trajectories was enhanced with puromycin in comparison to the CHX treatment (Figure S5D).

**Figure 5:**
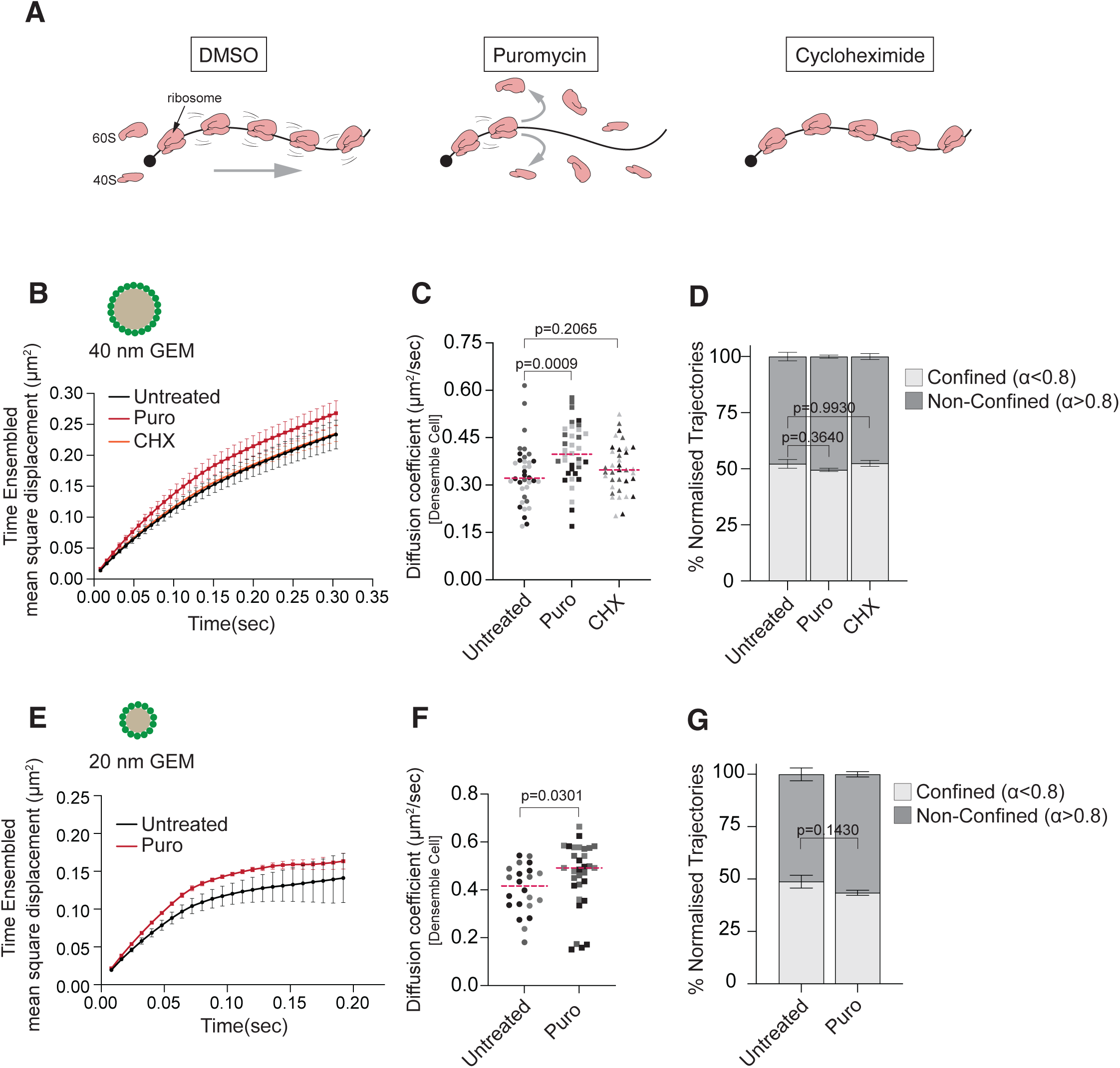
Mesoscale diffusion increases upon polysome disassembly in mammalian cells. A) Schematic illustration of translation inhibition with puromycin (Puro) and cycloheximide (CHX) for polysome disassembly and stabilization in mammalian cells. B) Time ensembled averaged mean square displacement of 40 nm GEMs upon treatment with Puro and CHX. Shown are means±SEM of three biological replicates performed on different days. C) Median diffusion coefficient of 40 nm GEMs upon treatment with Puro and CHX from three biological replicates performed on different days (median±SEM). Each data point represents the average diffusion coefficient of 40 nm GEMs in one cell. Different colors represent different days, and the dotted line represents the median value of the diffusion coefficients. Statistical comparison was performed using the Mann-Whitney test. Non-significant p value >0.05, significant p value <0.05. D) Polysome disassembly with Puro does not significantly change the confined trajectories of 40 nm GEMs in mammalian cells. Data shown here is from three independent experiments, and error bars are means±SEM. Statistical comparison was performed with two-way ANOVA with Šidák’s multiple comparison test. Non-significant p value >0.05, significant p value <0.05. E) Time ensembled averaged mean square displacement of 20 nm GEMs upon treatment with Puro. Shown are means±SEM of three biological replicates performed on different days. F) Median diffusion coefficient of 20 nm GEMs upon treatment with Puro from three biological replicates performed on different days (median±SEM). Each data point represents the average diffusion coefficient of 40 nm GEMs in one cell. Different colors represent different days, and the dotted line represents the median value of the diffusion coefficients. Statistical comparison was performed using the Mann-Whitney test. Non-significant p value >0.05, significant p value <0.05. G) Polysome disassembly with Puro does not significantly change the confined trajectories of 20 nm GEMs in mammalian cells. Data shown here is from three biological replicates and error bars represent means±SEM. Statistical comparison was performed with two-way ANOVA with Šidák’s multiple comparison test. Non-significant p value >0.05, significant p value <0.05.

We further asked whether smaller probes are also affected by polysome stability in mammalian cells. To test this, we generated 20 nm GEMs and transiently expressed them in HeLa cells. Upon acute treatment with puromycin, the mobility of these particles and their median diffusion coefficient (D_ensemble cell_) increased as well (Figure 5F). Similar to the experiments with 40 nm GEMs, we observed heterogeneity at an intercellular level (Figure 5G, S5F) and no significant change in the motion type of the particles based on our classification (Figure 5G). However, as observed for the 40 nm GEMs, a subtle increase in the confinement radius was also seen for the 20 nm GEMs upon polysome disassembly (Figure S5F). To examine whether these effects at the 20 nm scale are cell type-specific or more global, we also transiently expressed these particles in another cell line SkoV-3 and conducted analogous experiments. Consistent with our observations in HeLa cells, we observed comparable effects on MSDs and diffusion coefficients in the SkoV-3 cell line (Figure S5G&H). In conclusion, our results demonstrate that polysomes also affect the biophysical properties of mammalian cells at the mesoscale level.

## Discussion

The biophysical properties of a cell influence and control all cellular interactions and reactions. While an increasing number of studies have examined the rheology of the cytoplasm^8,27,60–64^, its regulation and impact on intracellular biochemistry are still poorly understood. Here, we employ single-particle tracking of endogenous mRNPs and GEMs of 20 nm and 40 nm size to demonstrate that the diffusion of particles in the size range of 20 to 200 nm depends on the translation status and the mRNA levels of cells. Thus, our study reveals that the higher-order organization of ribosomes into polysomes via mRNA is a critical regulator of the biophysical properties of yeast and mammalian cells at the mesoscale.

In this work, we examine exclusively the diffusivity of particles 20 nm and larger in proliferating cells. Our molecular simulation experiments would predict that macromolecules below ∼20 nm are no longer affected by the presence or absence of polysomes (Figure 3C). Thus, it should be expected that the diffusion of metabolites, individual proteins or even large spherical protein complexes up to a size of ∼ 2-3 MDa^65^ might not be affected by the higher-order organization of mRNAs and ribosomes into translating polysomes. Indeed, the magnitude of the effect of polysome disassembly on diffusion is reduced in 20 nm GEMs compared to 40 nm GEMs. However, we currently lack reliable tools to probe, at a single particle level, the cellular diffusion of macromolecules in size ranges below 20 nm. Therefore, our data do not clarify how polysomes and/or other smaller crowders influence the diffusion of protein complexes and other biomolecules that fall in this size range. Nevertheless, our results predict that multiple cellular processes that involve large macromolecular assemblies, such as cytoskeletal remodelling, movement and transport of vesicles and biomolecular condensates or signaling cascade complex interactions are influenced by the translational status of cells.

In a previous study, ribosome concentrations were modulated by either inhibiting the activity of Tor1 or by deleting the ribosome biogenesis factor *sfp1*^7^. However, inhibition of Tor1 and deletion of *SFP1* were previously also shown to lower translation levels and the number of polysomes in the cytoplasm^30,31^, and are also expected to lower the levels of several, very abundant groups of mRNAs coding for ribosomal proteins and ribosome biogenesis factors. Considering the results presented in our work, which reveal that the translation status and cellular mRNA levels directly affect cellular diffusivity at the mesoscale, these results need to be revisited. Interestingly, by perturbing ribosome numbers through deletion of the hibernation factor Stm1, we find that ribosome concentration alone cannot explain the change in mesoscale diffusivity. In *stm1Δ* cells, where ribosome numbers are decreased by 20-25% but translation is not significantly altered^45,46^, we do not observe any changes in diffusion of 40 nm GEM particles. This suggests crowding is dominated by polysomes and mRNA levels but not by free ribosomes in logarithmically dividing cells. However, in conditions where translation is significantly reduced, ribosome numbers might play a role in intracellular diffusion, which might explain some of the findings upon Tor1 inhibition or *SFP1* deletion^31,42^. This would be consistent with our simulations where we modulated ribosome numbers in the absence of polysomes (Figure 3C&S3E). Furthermore, we also see a small, albeit non-significant, increase in diffusion upon polysome disassembly with HIP in *stm1Δ* cells compared to wild-type cells. Upon *STM1* deletion, the reduction in ribosome number is also expected to be accompanied by a change in ribosome size distribution, with fewer 80S ribosomes and an increase in 60S and 40S subunits^46^, which could impact cellular crowding and colloidal pressure^3^. Understanding the contribution of the 60S and 40S crowding capabilities in translation-inhibited states will be intriguing to study in the future.

In addition to manipulating the translation status by drug treatment, we also lowered cytoplasmic mRNA levels, which led to an increase in the diffusion of GEM particles and thereby corroborated the significant role of polysomes as crowders in this size range. We accomplished this by acutely blocking mRNA export out of the nucleus through depletion of the mRNA export factor Dbp5. This leads to the nuclear retention of newly synthesized mRNAs, while the decay of previously exported mRNAs lowers cytoplasmic mRNA levels over time^29^. Due to the acute and relative short-term inhibition of mRNA export, minimal pleiotropic effects are expected. Indeed, we do not observe changes in the total amount of ribosomal proteins at the time point when we observe an increase in cytoplasmic diffusion. Thus, our Dbp5 depletion results show that mRNA levels in the cytoplasm control cytosolic diffusion. mRNA levels in the cytoplasm can be regulated by their biosynthesis, export and decay in the cell^66–68^, and it will be intriguing to investigate whether these pathways are used to control cytosolic diffusion.

Our experiments do not directly address whether ‘free’ mRNA outside of polysomes additionally impacts crowding and intracellular diffusion. RNA is a flexible polymer that in cells is thought to be organized in higher-order ribonucleoprotein complexes. Recent work has shown that mRNPs can adopt different conformations in cells, being more elongated when translated and more compact outside of polysomes^49^. In addition, secondary structure elements, RNA-RNA interactions in trans and various RNA-binding proteins are expected to influence the size and 3-dimensional organization of mRNP particles. Differences in mRNP compaction, organization and translation efficiency might explain the surprising finding that translating mRNAs that greatly vary in length display similar diffusive behavior in the cytoplasm (Figure 1G&J).

A recent paper by Xie *et al* concluded that the condensation of mRNA into P bodies controls the biophysical properties of yeast cells^69^. Interestingly, polysome disassembly with HIP results in mRNA condensation into P-bodies^57^. However, when we inhibit P-body assembly by depleting the DEAD-box ATPase Dhh1, no effect on GEM diffusion can be seen in the presence of HIP (Figure 4F). Thus, our results suggest that condensation of mRNAs into P-bodies does not have an effect on intracellular diffusion in yeast. Consistent with this, it was recently demonstrated that only a small percentage of the total mRNA population is sequestered into P-bodies in various stress conditions, and most untranslated mRNAs are localized outside microscopically visible P-bodies^70,71^. Currently, we cannot explain these differences in observations, but it is worth mentioning that Xie *et al*^69^ performed their experiments in glucose starvation whereas we analyzed diffusion in proliferating yeast cells. Xie *et al* also reported that G3BP1/2, two essential proteins for stress granule formation in mammalian cells^72^, are necessary for regulating intracellular diffusion upon polysome collapse in arsenite stress^69^. Therefore, the formation of mRNP granules and mRNA condensation might be a mechanism to control intracellular diffusion during stress but this might not be the case in proliferating yeast cells. Further work will be needed to better understand the role of polysome-free mRNA in cytoplasmic organization and crowding and to address how different mRNA states, e.g. compacted or condensed^49^, or the potential formation of small mRNP clusters outside of microscopically detectable P-bodies or stress granules impact intracellular diffusion. However, based on our results we can conclude that cytoplasmic mRNA levels impact the biophysical properties of the cytoplasm by organizing ribosomes into polysomes.

Our results demonstrate that the regulation of cytosolic diffusion by polysomes is conserved between yeast and mammalian cells. Similar to Xie *et al* we observed an increase in cytoplasmic diffusion upon polysome disassembly with puromycin. In contrast to yeast, we did not observe a significant change in the motion type in mammalian cells when we analyzed the percentage of the trajectories that were classified as confined or non-confined. One potential reason for this difference could be the lower ribosomal density in mammalian cells. However, both in yeast and mammalian cells we observed an increase in the radius of confinement upon polysome disassembly (Figure S2B&D and S5D&F). These results suggest that polysomes locally confine particle movement at a mesoscale. Whether this is brought about by individual mRNP-ribosome complexes or potentially also higher-order complexes that may form via RNA-RNA interactions or by co-translational complex assembly is unclear. Alongside polysomes, other subcellular compartments like TIS granules can form co-translation-dependent assemblies that might affect intracellular diffusion^73,74^. Moreover, the translation of large and abundant groups of mRNA occurs on the endoplasmic reticulum surface, which could also be a cause of local confinement. Further investigation is needed to determine whether such higher-order entities might affect intracellular diffusion and contribute to the observed changes in confinement in diffusion. Consistent with recent studies^7,8^, we also observe that diffusion in the cytoplasm is heterogeneous in both yeast and mammalian cells (Figure S1A&C, S2A&C, S5C). The source of this heterogeneity in the intracellular milieu is unclear but it will be interesting to understand how it correlates with intracellular compartmentalization and whether this heterogeneity contributes to the regulation of the cellular biochemistry.

## Material and Methods

### Strain generation

*Saccharomyces cerevisiae* strains were constructed using standard yeast genetic techniques either by transformation of a linearized plasmid or of a PCR amplification product with homology to the target site^75^.

### Yeast culture

Synthetic complete medium (SCD) containing 2% glucose was inoculated from individual colonies and grown overnight. Saturated cultures were diluted in fresh SCD at an OD of 0.15 and grown into the exponential phase (OD 0.6-0.8). All experiments were performed using cultures in the log phase (OD 0.6-0.8).

### Mammalian culture

Mammalian cells were maintained at 37°C and 5% CO_2_. HeLa and SkoV-3 cells were grown in DMEM, high glucose, GlutaMAX™ (Thermo Fischer Scientific,10566016) supplemented with 10% fetal bovine serum (Gibco™ 10500064) unless otherwise stated. All cell lines used in the experiments were passed for a maximum of 10 passages and were regularly checked for *Mycoplasma* contamination.

### Drug treatments

Drug treatments (final concentrations: Hippuristanol: 10 µM; Cycloheximide: 50 µg/ml) were added to 100 or 200 µl of cell suspension in a 1.5 ml microcentrifuge tube and mixed well by vortexing. 10 or 15 µl of cell suspension was plated onto a Concanavalin A coated well of a microscopy imaging plate. The plate was centrifuged at 100 x g for 2 min. Images were acquired between 15 and 30 min after the drug treatment. For mammalian cells, cells were treated with Hippuristanol (10 µM), Cycloheximide (10 µM) and puromycin (10 µM) and images were acquired between 20-30 min after the drug treatment.

### Live single-molecule RNA imaging

Yeast cells were transferred to a Concanavalin A-coated 8-well IBIDI dish and spun down for 1 min at 100 x g. Cells were imaged with a 150x objective (Olympus, AUPON150XOTIRF) and a laser power of 200-250 µW for GFP, 100-150 µW for mKate, using a 488 nm (Cobalt) and 594 nm (Coherent) lasers respectively. Images were acquired with 15 ms exposure (effective exposure 16.75 ms) for 1000 frames and a 201×201 pixel field of view using two EMCCD cameras (Andor iXon Ultra), which were synched (master/slave) for simultaneous imaging of two colours. The excitation light was filtered out before the objective with two single-notch filters (488 nm StopLine® single-notch filter and 594 nm StopLine® single-notch filter, Semrock). The emitted light was split to the respective cameras using a 580 nm edge BrightLine® single-edge imaging-flat dichroic beam splitter and filtered with 2 distinct emission filters (520/44 nm BrightLine® single-band bandpass filter for camera 1, 628/40 nm BrightLine® single-band bandpass filter for camera 2, Semrock). The signal-to-noise ratio of the GFP-tagged single particle images was improved using the Noise2Void algorithm^76^. Three stacks of 100 images were chosen to train the convolutional neural network, with a kernel size of 3, batch size of 128, and 100 training epochs.

### HILO Imaging of 40 and 20 nm GEMs

384-well plates (Matrical) were coated with concanavalin A for a few minutes. 10-15 ul of yeast cell suspension after respective treatments were plated in each well. The plate was centrifuged at 100 x g for 2 min and imaged at 30°C unless otherwise stated. Strains expressing 40 nm and 20 nm GEMs were imaged at 30-70% and 100% laser power, respectively, using 488 nm (80 mW) laser excitation on a Nikon N-STORM microscope in HILO (Highly Inclined and Laminated Optical) mode. Imaging was performed with an SR Apochromat TIRF 100X (NA=1.49) oil immersion objective and sCMOS camera: Hamamatsu Orca Flash 4 v3 (6.5 x 6.5 μm² pixel size). We use an additional demagnification lens system with a 2.4X de-magnifying power. The effective pixel size was 160nm (6.5um/100X*2.4X). Images were acquired at ∼100 frames/sec for 7 seconds using NIS Elements Advanced research software. Images were processed using Fiji software.

### Single particle localization and tracking

Tracking of particles (both mRNP and GEMs) was performed with the Track Mate^77^ plugin of FIJI. For the localization step, the LoG detector with sub-pixel localization enabled was used and for the tracking step, the Simple LAP tracker. For mRNPs: spot radius = 0.17 microns, quality threshold = 140.0. For yeast 40 nm GEMs: spot radius = 0.4 microns, quality threshold = 7.0-14.0. For yeast 20 nm GEMs: spot radius = 0.4 microns, quality threshold = 16.0. The majority of mRNAs in yeast are cytoplasmic^78,79^ therefore cell segmentation was not performed. For mammalian cells 40 nm GEMs: spot radius = 0.35 microns, quality threshold = 7.0, maximum number of frames to close the gap = 2. For mammalian cells 20 nm GEMs: spot radius = 0.4 microns, quality threshold = 35.0. For trajectories of mRNP and GEMs in yeast a minimum track length of 6 and 10 frames was used respectively. For trajectories of 20 and 40 nm GEMs in mammalian cells a minimum track length of 10 and 20 frames was used respectively.

### Trajectory classification

All generated trajectories were analyzed with a custom written Matlab(R2021) pipeline which uses the @msdanalyzer library^80^. The codes are available here: https://github.com/PabloAu/Single-Molecule-Tracking-Analysis/tree/master/V2. First, the Mean Squared Displacement (MSD) was computed for an individual trajectory and a motion type classification performed, to split the trajectories into confined and non-confined subgroups^81^. This classification was done by fitting a power law function to an individual MSD curve and obtaining the anomalous exponent coefficient α:

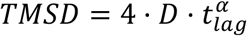

where D is the diffusion coefficient, t_lag_ is the time lag between the different time points of the track, and α is the anomalous exponent coefficient. Trajectories with α≤0.8 were considered as confined and trajectories with α>0.8 as non-confined. The percentage belonging to each population is shown in several figures.

### Trajectory analysis: diffusion coefficients

Diffusion coefficients were obtained from the trajectories in three different ways. First, in an individual fashion by fitting the first 4 points of each Time MSD curve with a linear function^82^.

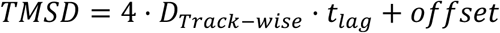

Second, in a population-based fashion, by fitting the first 4 points of the Time Ensemble MSD (TEMSD) curves with a linear function:

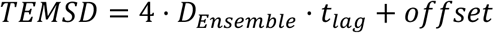

This loses information about trajectory-to-trajectory variation, but it is more accurate since the number of jumps to compute the TEMSD curves is much higher^81^. Both the track-wise and ensemble diffusion coefficients (D_ensemble_) are shown across the different figures. Third, an ensemble cell-by-cell analysis was performed, where all the trajectories from single cells were used to compute the TEMSD. The resulting diffusion coefficients are referred to as D_ensemble cell_

### Trajectory analysis: radius of confinement

The radius of confinement was obtained only for the subgroup of confined trajectories. It quantifies the degree of confinement by estimating the radius of the potential volume explored by the particle. It was measured by fitting a circle-confined diffusion model to the TEMSD (ensemble of all trajectories)^83^

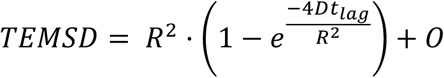

where R is the radius of confinement and D is the diffusion coefficient at short timescales. O is an offset value that comes from the localization precision limit inherent to localization-based microscopy methods.

### Auxin degron depletion

Auxin-inducible degradation of proteins was induced by treatment with 0.5 mM indole-3-acetic acid (IAA) (Sigma, I2886-5G, CAS: 87-51-4) and 4 μM phytic acid dipotassium salt (Sigma, 5681, CAS: 129832-03-7) for the indicated time points.

### Western blot

Logarithmically growing cells were collected and normalized according to O.D. The pellet was lysed with 0.1 M NaOH for 15 min. Samples were spun down at 4000 x g for 5 min and the pellet was resuspended in 60 µl of 1x SDS loading buffer. Normalized lysates were separated in 12 % acrylamide SDS-PAGE gels. Blotting of proteins was done with a Biorad semi-dry system in TBE buffer (pH=8.3) at 25V and 1 Amp followed by a Ponceau-S staining to confirm transfer. The following primary antibodies were used at the indicated dilutions: rabbit-anti-α-S5 (1:4000)^84^, rabbit-anti-α-L35 (1:4000)^84^, rabbit-anti-Hexokinase (1:3000) (US Biological, Cat no. 169073), mouse-anti-V5 (1:5000) (Sigma-Aldrich). IRDye® 800CW goat anti-rabbit (1:10000) (LI-COR Biosciences GmbH, Cat no. P/N 926-32211), Alexa 680 goat anti-mouse (Thermo Fisher, Cat no. A-21057) were used as secondary antibodies for detection.

### Lattice light sheet microscopy

3×10^4^ cells were plated on an Ibidi µ-slide 8-well glass bottom plate (Cat. No. 80827). Cells were grown in DMEM, high glucose, GlutaMAX™ (Thermo Fischer Scientific,10566016) for 24 hrs and washed with 1x PBS to remove the medium. Imaging medium (FluoroBrite™ DMEM (A1896701), 10% fetal bovine serum (Gibco™ 10500064), sodium pyruvate and glutamine) was added to the wells before imaging. Cell lines expressing GEMs were imaged using 488 nm laser excitation with 90% laser power, at 5 ms exposure time with light sheet dimensions (100X1800 μm) with a Zeiss lattice light sheet microscope. Imaging was performed with a 48X/1.0NA detection objective and Hamamatsu Fusion sCMOS camera. Images were acquired at ∼119 frames/sec for 5-6 sec using ZEN blue software. Images were processed using Fiji software.

### Transient transfection of 20 and 40 nm GEMs

For transient transfection, HeLa or Skov-3 cells were seeded as 60%-70% confluency in an Ibidi µ-slide 8-well glass bottom plate (Cat. no. 80827) on the day before transfection and were transfected with 700 ng of plasmid DNA (pKW4637 or pKW5385) using Lipofectamine™ 2000 Transfection Reagent (11668019) as per manufacturer instructions. 16-18 h post-transfection, fresh DMEM, high glucose, GlutaMAX™ was used to replace the transfection mixture. Cells were allowed to recover for 24 hrs, and imaging was performed.

### Brownian dynamics simulations

All Brownian dynamics (BD) simulations were performed with GROMACS 2022. All particles are represented as soft spheres with sizes relative to the size of the ribosome, which is taken equal to 1σ. The size of the ribosome is in the range 25-30 nm, thus 20nm GEMs and 40nm GEMs particles have reduced diameters in the interval of 0.666-0.8σ and 1.333-1.6σ respectively. The short-range (SR) interaction potential is composed of the repulsive part of the Lennard-Jones potential, of the form

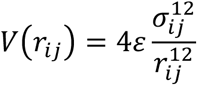

Where r_ij_ is the distance between the particles, σ_ij_ is the arithmetic average of the diameters of the two particles, and ε is the energy scale equal to 1/4 k_B_T, where k_B_ is the Boltzmann constant, and T is the temperature in Kelvin. The energy scale has been chosen to have a repulsive potential of 1 k_B_T at distance σ_ij_. SR interactions are truncated at 2.5σ for simulations with probe particles of size smaller than 2σ, in simulations with probe particles of size 2σ, 3σ, and 4σ, cut-off distances were respectively at 5σ, 6σ, and 7σ. Representative potential energy profiles are shown in Figure S3B. Polysomes are represented as spring-bead polymers of ribosomes. To estimate the parameters for the harmonic potential between consecutive ribosome particles in a polysome we started from the following hypotheses: i) mRNA can be approximated as an ideal chain of segments with a persistence length of 1.5 nm and Kuhn length (l) of 3 nm, ii) the average distance in RNA between ribosomes is 154 nt, and iii) the spacing between bases is 2 bases/nm. Thus, the average distance between consecutive ribosomes is ∼75 nm or 25 Kuhn lengths. The distribution of end-to-end distances for an ideal chain is:

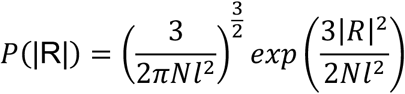

With an average value of end-to-end distance equal to:

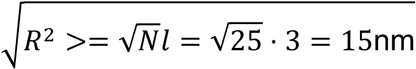

Considering 1σ (the size of the ribosome) equal to 30nm, the average end-to-end distance is 0.5σ. To consider the steric exclusion between ribosomes, an additional σ is imposed to the distance, giving the equilibrium distance for the bonded potential R_0_=1.5σ (Figure S3C). The potential energy associated with the end-to-end distance distribution is (Figure S3D):

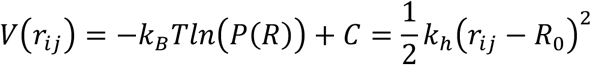

Where C is an immaterial constant added to have min(V)=0. From this equality we obtain kh=29.939 kJ/mol/σ^2^. The SR interaction between consecutive ribosomes is not excluded. The resulting potential between consecutive beads in a polysome is shown in Figure S3E.

To define the length of polysomes, we remapped the distribution of lengths of mRNA molecules obtained from mRNA sequencing to a distribution of number of ribosomes per mRNA molecule, assuming an average of ∼5 ribosomes per mRNA molecule. We fitted the distribution of the number of ribosomes on RNA molecules to a lognormal distribution (Figure S3F) and we sampled the fitted distribution to define the polysome composition of each simulation. Each replica has a different composition of polysomes to avoid bias due to specific compositions. Representative compositions and comparison with the objective distribution are reported in Figure S3G, note that lengths equal to 1 are possible.

In each simulation, there are 1000 ribosome particles, divided between polysomes and free ribosomes according to the simulated fraction of ribosomes not involved in polysome formation, and 10 probe particles. Concentrations (c) are referred to the physiological value of 13000 ribosomes/µm^3^ (12.17 μM), which is taken as c=1 in reduced units. The side of the cubic simulation box at c=1 is 14.176σ. When changing the concentration of the system, the number of particles is kept constant, and the size of the box is varied accordingly. In simulations mimicking *STM1* deletion, the concentration of the system was set to c=0.75.

Initial configurations were generated by placing the particles (ribosomes and GEMs) and polysomes in random positions and orientations using the gmx insert-molecules tool in GROMACS. Polysome chains are inserted in an extended configuration with a distance between ribosomes equal to the equilibrium value. Systems were relaxed with 1000 steps of minimization using the steepest descent algorithm to remove steric overlaps between the particles that could arise from the random insertion. For the integration of the equations of motion, we set the friction coefficient to 10 Da/t* for all particles, where t* is the simulation time unit, while the timestep has been set to 0.0005 t*, as it gives the correct temperature reported by gmx energy tool, as suggested in the GROMACS user manual. In Figure S3A we report evaluated temperatures for different timestep values. Each simulation was run for 200 million steps, saving conformations of all particles every 2,000 steps for further analysis.

The mean square displacement (MSD) of probes was evaluated using gmx msd tool, after unwrapping the trajectories of the probe particles using the –pbc nojump flag in gmx trjconv to have continuous movement of all particles. All saved frames were used and the maximum time delta between frames to calculate MSDs (-maxtau flag) was set to 20,000 t*. The diffusion coefficient D for each particle was then evaluated by fitting the MSD(t) trace using:

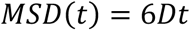

For each system, 10 replicas were run. Reported values are an average of replicates and errors are estimated as standard error of the mean with n=10.

---

**Supplementary table 1.** (List of strains)

| Strain name | Genotype | Figure | Reference |
| --- | --- | --- | --- |
| KWY6353 | *MAT A his3Δ1 leu2Δ0 ura3Δ0 NDC1::NCD1-yomKate2-URA3 GFA1::GFA1-24xPP7stemloops pMET25-PP7CP-yEGFP::LEU2* | Figure 1F | This study, base strain is a gift from Montpetit lab |
| KWY6352 | *MAT A his3Δ1 leu2Δ0 ura3Δ0 NDC1::NCD1-yomKate2-URA3 TRA1::TRA1-24xPP7stemloops pMET25-PP7CP-yEGFP::LEU2* | Figure 1 | This study, base strain is a gift from Montpetit lab |
| KWY7664 | *his3D1/his3Δ1 ura3Δ0/ura3Δ0 MDN1/MDN1-24PP7sl-KANleu2Δ0/LEU2::MET25(pro)-PP7CP-yEGFP NDC1/NDC1-mKate::HIS* | Figure 1 | This study |
| KWY6116 | *MAT A his3Δ1 leu2Δ0 ura3Δ0 NDC1::NCD1-yomKate2-URA3GFA1::GFA1-24xPP7stemloops::loxP-kanR-loxP pMET25-PP7CP-yEGFP::LEU2* | Figure 1B | This study |
| KWY8913 | *MAT A his3-11,15 ura3-1 trp1-1 LEU2::pINO4-PfV-GS-Sapphire (pKW4635)*<br>40nM GEM integrated into the LEU2 locus | Figure 2 | This study |
| KWY11346 | *MAT A his3-11,15 ura3-1 trp1-1 pHIS-Aq_LumSyn-Sapphire (pKW4636)*<br>20nM GEM integrated into the URA3 locus | Figure 2 | This study |
| KWY11835 | *MAT A W303 mat A stm1::NATMX6 LEU2::pINO4-PfV-GS-Sapphire (pKW4635)*<br>40nM GEM integrated into the LEU2 locus | Figure 2 | This study |
| KWY11251 | *DBP5-3V5-IAA7:: KANMX LEU2::pINO4-PfV-GS-Sapphire (pKW4635)* | Figure 4 | This study |
| KWY11253 | *DBP5-3V5-IAA7:: KANMX* | Figure 4 | This study |
|  | *LEU2::pINO4-PfV-GS-Sapphire* (pKW4635)<br>*pNH603-pGPD1-osTIR::HIS* (pKW2874) |  |  |
| KWY11651 | *MAT A dhh1::DHH1-3V5-IAA7::KANMX*<br>PKW4635 (37 nm GEMs) integrated into LEU2 Locus | Figure 4 | This study |
| KWY11652 | *MAT A dhh1::DHH1-3V5-IAA7::KANMX*<br>PKW4635 (37 nm GEMs) integrated into LEU2 Locus<br>PKW2874 (OsTIR) integrated into HIS3 Locus | Figure 4 | This study |

**Supplementary Table 2.** (Plasmid List)

| Plasmid name | Description | Figure | Reference |
| --- | --- | --- | --- |
| pKW4635 | pLH497 | Figure 1 | <sup>7</sup> |
| pKW4636 | pLH1144 | Figure 1 | <sup>7</sup> |
| pKW4637 | pLH611 | Figure 5 | <sup>7</sup> |
| pKW5385 | pcDNA3.1-PfV-Sapphire 20nM GEM (AMP) | Figure 5 | This study |

## Author contributions

Conceptualization: V.G and K.W.

Methodology: V.G, P.G.G, A.D, S.H, and K.W.

Investigation: V.G, S.H, M.P, P.G.G, D.M, A.D.

Formal analysis: V.G, P.G.G, M.P and A.D.

Writing-Original draft: V.G, and K.W.

Writing-Review & Editing: V.G, S.H, M.P, P.G.G, S.K, and K.W.

Funding acquisition: K.W.

Resources: V.G, K.W.

Supervision: V.G, A.B. and K.W.

## Conflict of interest

The authors declare no competing interests.

## Acknowledgements

The authors are grateful to Sébastien Huet, Yoav Shechtman, Thomas Michaels, Sandeep Choubey and Sider Penkov for their comments on the manuscript. We thank members of the Weis lab for suggestions on the manuscript. We acknowledge Roberta Mancini for strain generation and Matthias Neeracher for technical assistance. We thank Tobias Schwarz, Joachim Hehl, Dorothea Pinotsi, Sungsik Lee and Justine Kusch from the ETHZ ScopeM facility for their technical assistance and help with microscopy. We thank the Kapoor and Holt labs for providing the HeLa cell line expressing 40 nm GEMs, the Zampieri lab for the SkoV-3 cell line, the Montpetit and Singer labs for providing yeast strains and Michaela Oborska and Vikram Panse for providing the RPS5A and RPL35A antibodies. We are grateful to Eliana Bianco and Matthias Peter for providing the *STM1* deletion strain and for sharing the results before publication. We are indebted to Junichi Tanaka for the kind gift of Hippuristanol. P.G.G. acknowledges support from an ETH postdoctoral fellowship (22-1 FEL-06). This work was further supported by an EMBO long-term fellowship (ALTF 290-2014, EMBOCOFUND2012, GA-2012-600394 to S.H.) and by grants from the Swiss National Science Foundation (SNF CRSII5_193740 to A.B. and K.W., and 310030_208213 and TMAG-3_209354 to K.W.).

---

**Supplementary Figure 1:**
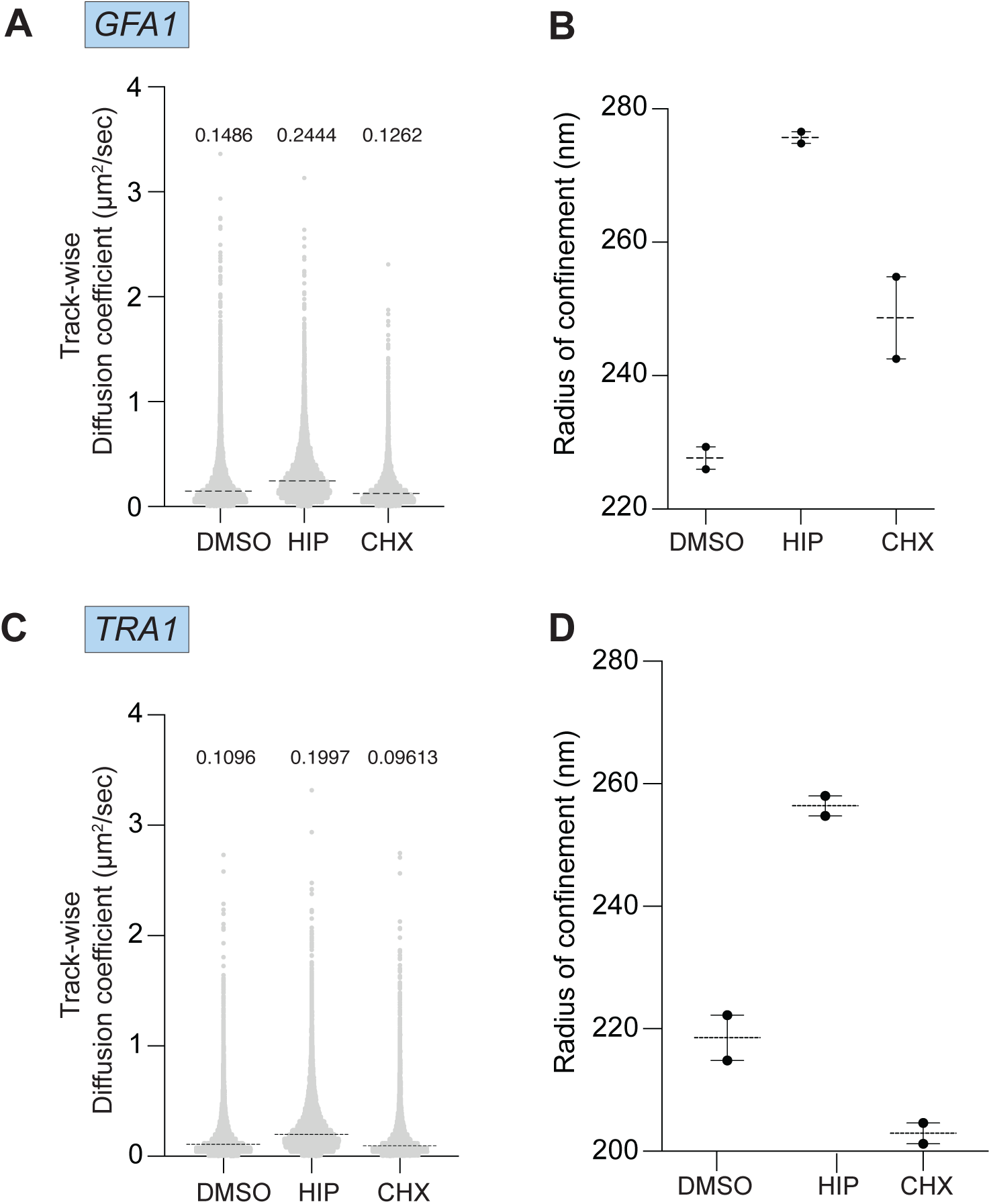
Polysome disassembly with HIP increases the confinement radius of mRNPs. A) Track-wise diffusion coefficients of GFA1 mRNP from two biological replicates. Each data point represented here is the diffusion coefficient of a single track (n) generated by mRNPs. DMSO (n = 7658), HIP (n = 8175), CHX (n = 6652). The dotted line and the number above represent the median value of the diffusion coefficients. B) Confinement radius of GFA1 mRNP increases upon polysome disassembly with HIP. Shown is the mean radius of confinement upon polysome disassembly with HIP from two biological replicates. C) Track-wise diffusion coefficients of TRA1 mRNP from two biological replicates. Each data point represented here is the diffusion coefficient of a single track (n) generated by mRNPs. DMSO (n = 8274), HIP (n = 10658), CHX (n = 9268). The dotted line and the number above represent the median value of the diffusion coefficients. D) Confinement radius of TRA1 mRNP increases upon polysome disassembly with HIP. Shown is the mean radius of confinement upon polysome disassembly with HIP from two biological replicates.

**Supplementary Figure 2:**
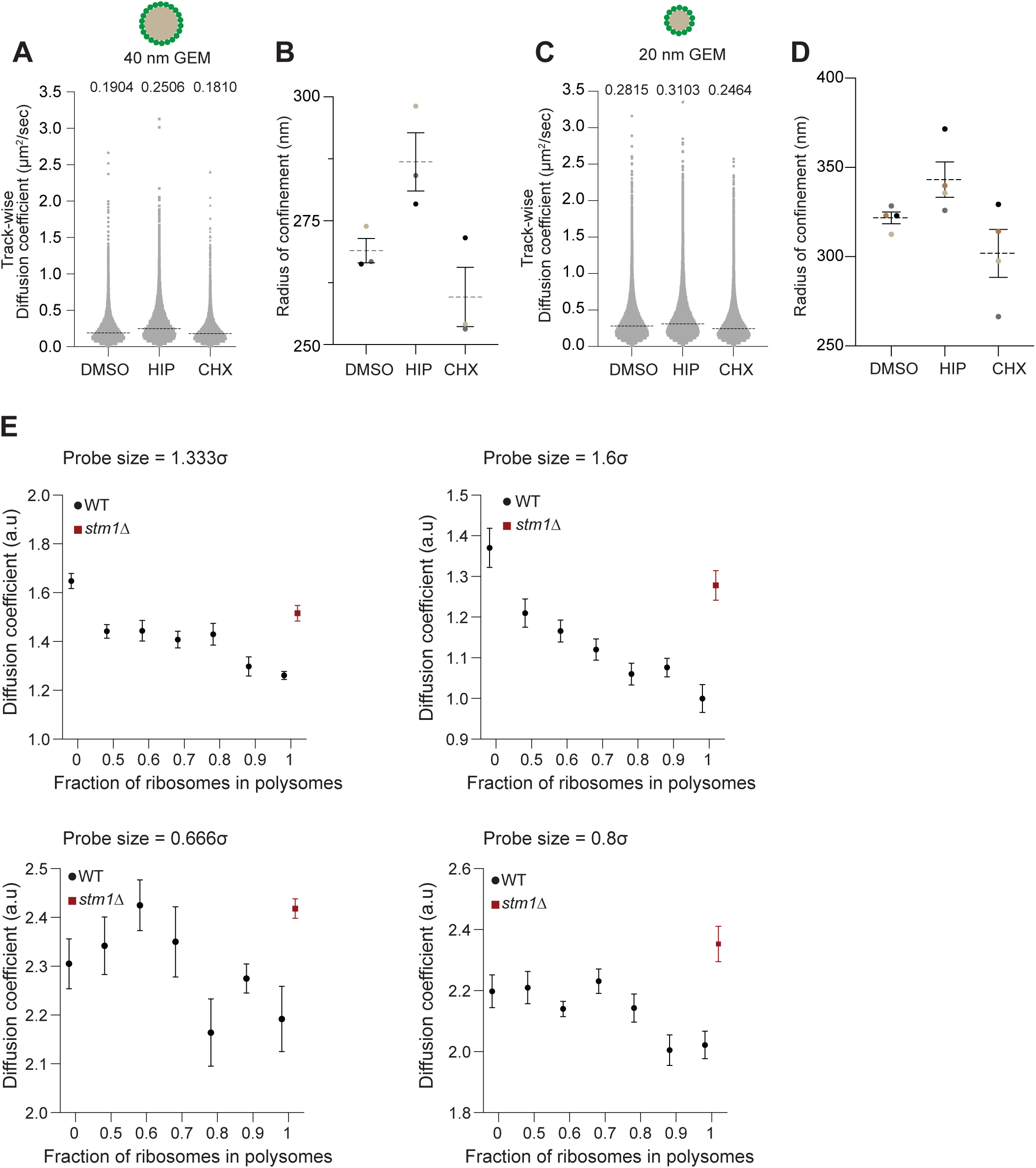
HIP treatment increases confinement radius of 40 and 20 nm GEMs in budding yeast. A) Track-wise diffusion coefficients of 40 nm GEMs in yeast from three biological replicates. Each data point represented here is the diffusion coefficient of a single track generated by 40 nm GEMs. DMSO (n = 12,756), HIP (n = 13,839), CHX (n =13510). The dotted line and the number above represent the median value of the diffusion coefficients. B) Confinement radius of 40 nm GEMs increases upon polysome disassembly. Shown is the mean±SEM radius of confinement of 40 nm GEMs upon polysome disassembly with HIP from three biological replicates. C) Track-wise diffusion coefficients of 20 nm GEMs in yeast from four biological replicates. Each data point represented here is the diffusion coefficient of a single track generated by 20 nm GEMs. DMSO (n = 41,089), HIP (n = 49,315), CHX (n =41,499). The dotted line and the number above represent the median value of the diffusion coefficients. D) Confinement radius of 20 nm GEMs increases upon polysome disassembly. Shown is the mean±SEM radius of confinement of 20 nm GEMs upon polysome disassembly with HIP from four biological replicates. E) Diffusion coefficients from Brownian dynamics simulations for probes of size ranges corresponding to 40nm (top row) and 20nm (bottom row) GEM particles. Black points indicate values from simulations at physiological concentration of ribosomes at varying fractions of ribosomes involved in polysome formation. Red squares indicate values from simulations mimicking the effect of STM1 deletion, where all ribosomes are involved in polysome formation, but the total concentration is 75% of the physiological value. Values of diffusion coefficients are average values over 10 independent simulations, error bars indicate SEM.

**Supplementary Figure 3:**
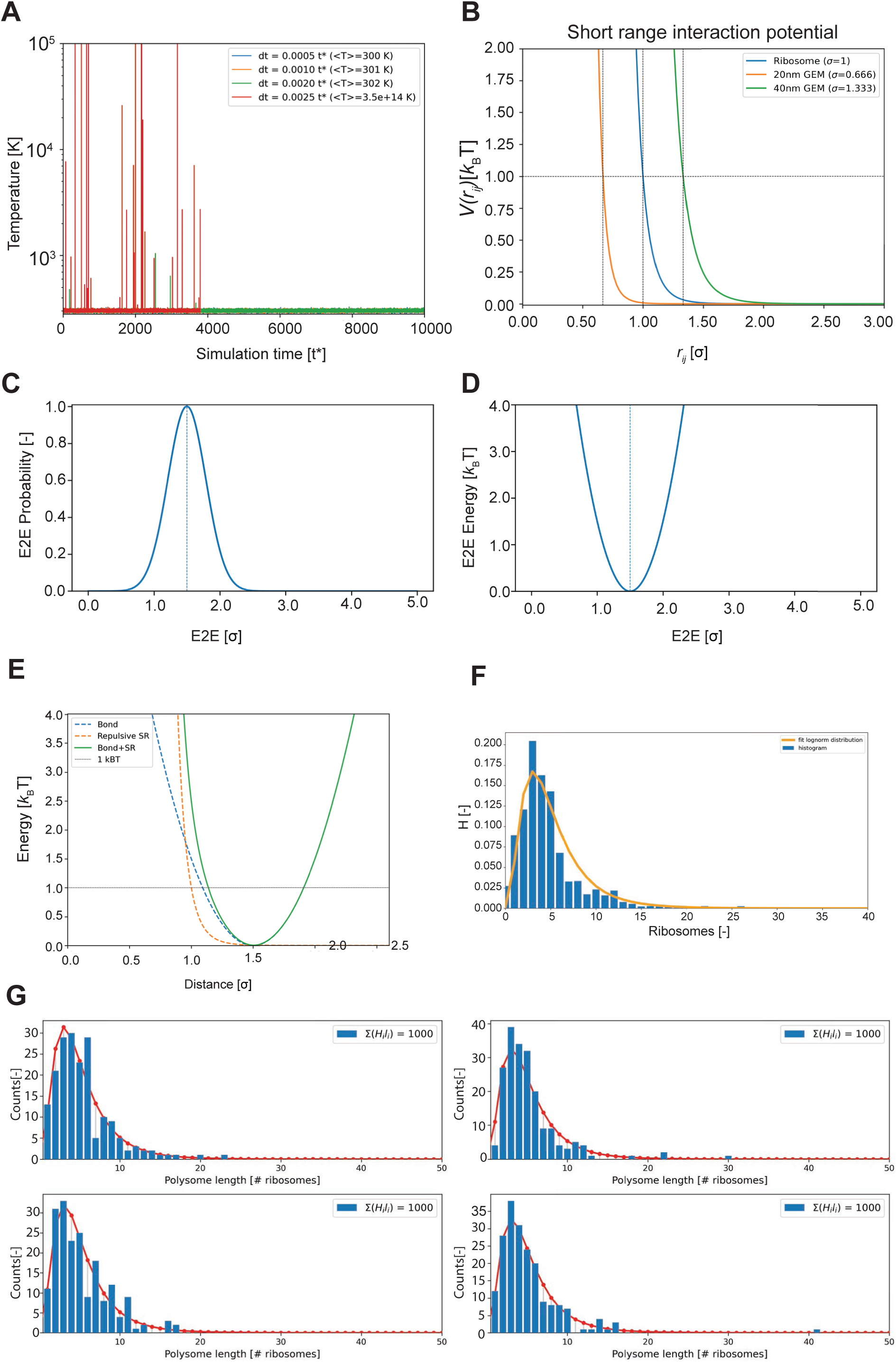
Parametrization of the molecular model of polysome organization. (A) Estimated temperatures from Brownian dynamics simulations at different time steps for the integration of the equations of motion. Time steps of 0.0005 t* allow for an accurate control of the temperature of the systems, while larger time steps introduce integration errors resulting in an inaccurate control of the temperature. (B) Representative profiles for the excluded volume interaction between the particles in the system. (C, D) Probability distribution of the distance between the centers of two consecutive ribosomes, assuming that they are connected via an ideal chain with an average distance that is 1.5 times the size of a ribosome (C) and the associated free energy profile (D). (E) Potential energy for the interaction between two consecutive ribosomes in a polysome: dashed lines represent the components of the interaction, namely the bonded harmonic potential and the excluded volume short-range interaction; solid line is the sum of the two components. The horizontal indicates the 1 kBT energy level. (F) The distribution of the number of ribosomes on mRNA molecules from mRNA sequencing data (bars) has been fitted to a lognormal distribution (line) assuming an average of ∼5 ribosomes per mRNA molecule. The obtained distribution has been then used to sample populations of polysome lengths. (G) Representative samples (bars) extracted from the fitted distribution (lines) indicate that the total number of ribosomes in the system is equal to 1000.

**Supplementary Figure 4:**
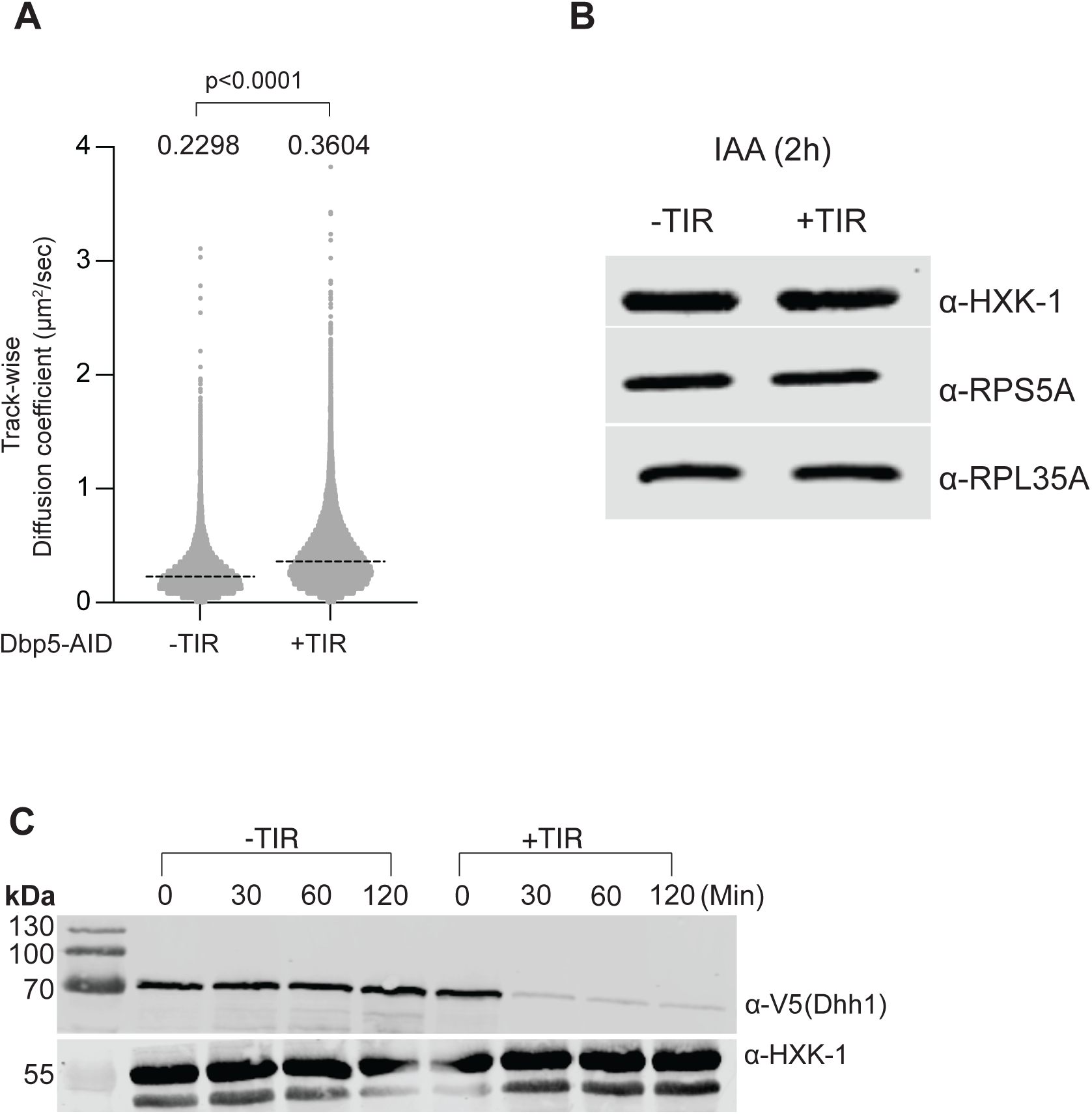
Acute depletion of Dbp5 does not affect total ribosomal amounts. A) Track-wise diffusion coefficients of 40 nm GEMs upon Dbp5 depletion. Statistical comparison was performed using the Kolmogorov-Smirnov test. Non-significant p value >0.05, significant p value <0.05. The dotted line and the number above represent the median value of the diffusion coefficients. B) Western blot of ribosomal proteins of the large subunit (Rpl35A) and small subunit (Rps5A) following Dbp5 depletion, with hexokinase (Hxk-1) as a loading control. Data shown here is a representative image of three biological replicates. C) Acute depletion of Dhh1-AID. Data shown here is from one experiment.

**Supplementary Figure 5:**
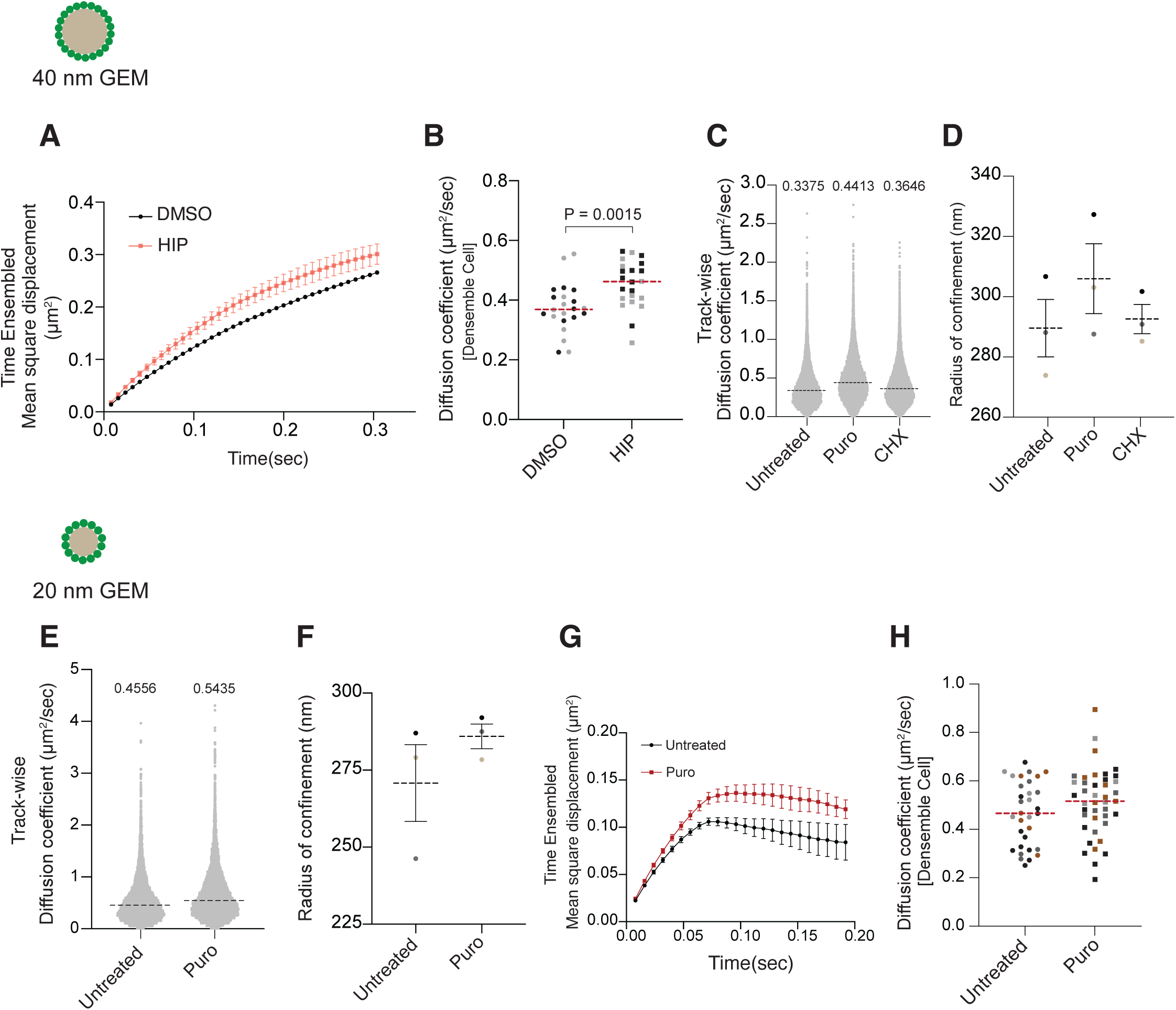
Hippuristanol and puromycin treatment enhances diffusion of 40 and 20 nm GEMs in mammalian cells. A) Time ensembled averaged mean square displacement of 40 nm GEMs upon treatment with HIP. Shown are means±SEM of two biological replicates performed on different days. B) Median diffusion coefficient of 40 nm GEMs upon treatment with HIP. Each data point represents the average diffusion coefficient of 40 nm GEMs in one cell. Different colors represent different days, and the dotted line represents the median value of the diffusion coefficients. Statistical comparison was performed using the Mann-Whitney test. Non-significant p value >0.05, significant p value <0.05. C) Track-wise diffusion coefficients of 40 nm GEMs in mammalian cells upon treatment with Puro and CHX from three biological replicates. Each data point represented here is the diffusion coefficient of a single track generated by 40 nm GEMs. The dotted line represents the median value of the diffusion coefficients. Untreated (n = 18874), Puro (n = 12304), CHX (n =18060). D) Confinement radius of 40 nm GEMs increases upon polysome disassembly with Puro. Shown are the median radius of confinement of 40 nm GEMs upon polysome disassembly with HIP from three biological replicates. E) Track-wise diffusion coefficients of 20 nm GEMs in mammalian cells upon treatment with Puro from three biological replicates. Each data point represented here is the diffusion coefficient of a single track generated by 20 nm GEMs. The dotted line represents the median value of the diffusion coefficients. Untreated (n = 8699), Puro (n = 7897). F) Confinement radius of 20 nm GEMs increases upon polysome disassembly with puromycin. Shown are the median radius of confinement of 20 nm GEMs upon polysome disassembly with Puro from three biological replicates. G) Time ensembled averaged mean square displacement of 20 nm GEMs upon treatment with Puro in SkoV-3 cells. Shown are means±SEM of three biological replicates performed on different days. H) Median diffusion coefficient of 20 nm GEMs upon treatment with puromycin. Each data point represents the average diffusion coefficient of 20 nm GEMs in one cell. Each data point represents the average diffusion coefficient of 20 nm GEMs in one cell. Different colors represent different days, and the dotted line represents the median value of the diffusion coefficients.

## Notes

### Competing Interest Statement

The authors have declared no competing interest.

